# Understanding the biomechanical and physiological responses to Advanced Footwear Technology in well-trained male and female runners

**DOI:** 10.64898/2026.06.19.732297

**Authors:** Yumna Albertus, David Leith, Oloff Bergh, Zachary B. Barrons, Nicholas Tam

**Affiliations:** Health through Physical Activity and Sport Research Centre, Faculty of Health Sciences, University of Cape Town, South Africa; Carnegie School of Sport, Leeds Beckett University, United Kingdom; Neuromechanics Unit, Central Analytical Facility, Stellenbosch University, Cape Town, South Africa; Sports Science Laboratory, On AG, Switzerland

**Author notes:** Address for correspondence: Yumna Albertus.

## Abstract

Advanced footwear technology (AFT) has transformed competitive running, yet individual and sex-specific responses to different AFT models remain unclear, particularly near race pace. This study examined running economy (RE) and gait biomechanics in response to three top-tier AFT models (Shoe A: adidas Pro Evo 2; Shoe B: Nike Alphafly 3; Shoe C: On CloudBoom Strike 2) in 14 male and 12 female well-trained runners at sex-specific submaximal speeds (16 and 14 km·h⁻¹). RE, spatiotemporal, and joint kinematic/kinetic data were collected via indirect calorimetry, accelerometry, and three-dimensional motion capture with force platforms. RE was significantly lower in Shoe C than Shoe A (males: 2.1%; females: 1.4%) and Shoe B (males: 1.9%; females: 0.9%), with 73% of runners responding favourably to Shoe C, a more consistent response than previously reported. Despite being lightest, Shoe A produced the poorest RE, challenging conventional mass-economy assumptions. Biomechanically, Shoe C elicited greater impact magnitude, lower ankle quasi-stiffness, and greater ankle angular velocity during early stance. Female runners showed smaller RE improvements, potentially related to lower running velocity and body mass limiting midsole engagement. The most efficient AFT enabled these well-trained runners to be more spring-like through tolerating higher forces and faster angular velocities without greater demand on metabolic cost.

## Introduction

Advanced Footwear Technology (AFT), characterized by longitudinal bending stiffness (LBS) moderators, high-resiliency foams, and pronounced sagittal-plane rocker profiles, has fundamentally altered competitive running by significantly improving performance across a wide range of distances and disciplines.^1^ On race day, it’s no longer a question of whether performance-oriented runners are wearing AFT, it’s a question of which AFT provides the best gain. While AFT is often discussed as a homogenous category, the current market has evolved into a diverse landscape of competing designs. From proprietary foams to varied plate geometries and rocker profiles, modern AFT takes many forms. This raises the practical question: how should an athlete choose which specific AFT model to wear on race day?

Initial investigations into components that later became defining features of AFT, such as longitudinal bending stiffness, introduced the concept of metabolic ‘responders’ versus ‘non-responders suggesting that improvements in running economy were highly individual. This phenomenon may indicate that the metabolic benefits of these interventions are sensitive to individual biomechanical interactions. For example, Madden et al. (2015) observed that while increasing forefoot bending stiffness did not significantly impact running economy at the group level, 55% of participants experienced individual metabolic improvements.^2^ These responders exhibited unique kinematic adaptations, most notably a decrease in peak ankle angular velocity. The authors posited that this reduction in velocity may optimize the force-velocity relationship of the plantar flexors, reducing the metabolic cost of muscular contraction by allowing the muscles to operate at slower, more efficient shortening velocities. Similar inter-individual variability has been observed in modern AFT comparisons; Hunter et al. (2019) found that the running economy improvements associated with a single AFT model spanned from 0% to 6.2%.^3^ More recent evidence suggests a more “universal” response. Kuzmeski et al (2026) found in a comparison of four AFT that all fifteen participants positively responded to a single model.^4^ Of those fifteen participants however only four were female, precluding researchers from determining whether sex specific differences in response may exist.

While both men’s and women’s records have been repeatedly shattered, the impact has been asymmetric.^5^ Women’s road race world records have improved by an average of 3.7% whereas men’s records have only improved by 1.5%.^6^ There is limited AFT research that has adequately represented the female sex. The evidence that does exist suggests that the biomechanical adjustments distinguishing high responders from low or non-responders may be sex-specific.^7^ However, due to inadequate power or methodological flaws strong conclusions could not be drawn.^7,8^

Lastly, most previous research have reported changes in running economy at similar relative intensities rather than absolute velocities, making it difficult to distinguish whether the observed biomechanical responses and subsequent mechanisms are influenced by differences in running speed as well as the AFT footwear itself.^9^ This is often limited by the availability of similarly well-trained high performing athletes capable of running sub-maximally at high-velocities.

Therefore, the purpose of this study was i) to examine the running economy and biomechanical responses to three current top tier AFT models in well-trained male and female runners at submaximal running speeds. We hypothesised that within each sex, the most economical AFT model would vary among individuals. Furthermore, we hypothesised that the biomechanical adaptations associated with improved RE would differ between shoe conditions for both male and female runners.

## Methods

### Overview of Testing

Twenty-six trained runners, 14 male and 12 female runners participated in this study and were able to run 10 km in <35 and <40 minutes respectively (Table 1). All runners were free of injury 6 months prior to participating in this study. Before participating, all runners signed an informed consent, while ethical approval for the study was granted by the human research ethics committee of the University of Cape Town.

**Table 1:** Descriptive characteristics of male and female participants (n=26)

|  | Age (yr) | Height (cm) | Body Mass (kg) | 10km Personal Best |
| --- | --- | --- | --- | --- |
| **Males (n = 14)** | 32 ± 7 | 175.4 ± 5.4 | 64.8 ± 6.4 | 32:22 ± 4:48 |
| **Females (n = 12)** | 30 ± 5 | 165.7 ± 6.3 | 53.1 ± 3.3 | 38:02 ± 3:13 |

Data are presented as mean ± standard deviations

### Footwear

Three AFT models were selected in this study namely, adidas Pro Evo 2 (Shoe A), Nike Alphafly 3 (Shoe B), On CloudBoom Strike 2 (Shoe C). These AFTs were characterised to understand their mechanical properties (Figure 1, Table 2).

**Figure 1:**
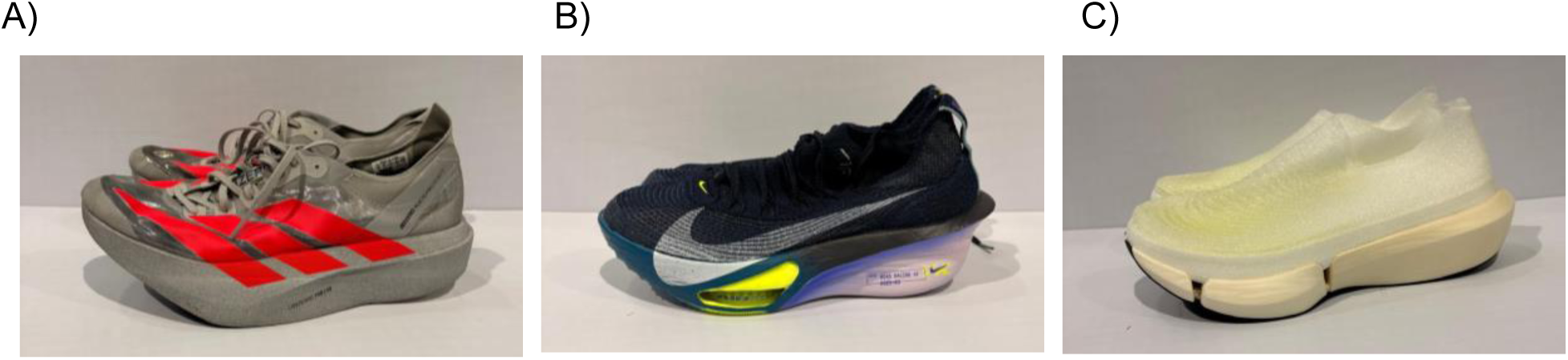
AFT Models (A) adidas Pro Evo 2 (Shoe A), (B) Nike Alphafly 3 (Shoe B) and (C) On CloudBoom Strike 2 (Shoe C)

**Table 2:** Mechanical properties of the different footwear conditions in the size US10.

|  | Shoe A<br>(adidas Pro Evo 2) | Shoe B<br>(Nike Alphafly 3) | Shoe C<br>(On CloudBoom Strike 2) |
| --- | --- | --- | --- |
| **Mass (g)** | 139 | 218 | 174 |
| **Midsole thickness (mm)** |  |  |  |
| Rearfoot | 39 | 38.1 | 40 |
| Forefoot | 36 | 29.6 | 36 |
| **Three-point bending** |  |  |  |
| LBS ($\text{N} \cdot \text{mm}^{-1}$) | 20.4 | 21.38 | 17.4 |
| **Rearfoot** |  |  |  |
| Deformation (mm) | 25.2 | 24.9 | 25 |
| Energy Loss (J) | 1.8 | 1.42 | 1.29 |
| Energy Return (J) | 11.97 | 11.63 | 11.6 |
| **Forefoot** |  |  |  |
| Deformation (mm) | 20.38 | 21.38 | 22.6 |
| Energy Loss (J) | 1.58 | 1.62 | 1.63 |
| Energy Return (J) | 9.8 | 11.45 | 11.3 |

Shoes were mechanically tested with compression and three-point bending tests. Compression testing was performed on both the heel and the forefoot. The 50-mm diameter impactor was positioned at the heel/forefoot and a preload of 25 N was applied. The forefoot was loaded with a force of 1800 N, while the heel was loaded with 1500 N, both at a frequency of 2 Hz. Compression stiffness was calculated as the maximum force divided by the total displacement for each cycle. The average of 10 compression cycles was used to compute the compression stiffness of the midsole. For the three-point bending test to determine LBS, the shoe midsole was displaced vertically by 15 mm, with the supporting pins placed 140 mm apart. The displacement speed was set to 15 mm·s^−1^, with each cycle consisting of 1 s of downward displacement and 1 s of upward return for six cycles. Stiffness values were first averaged over the linear region of the force-displacement curve, defined as 80% – 90% displacement of each loading cycle, and then averaged across all six cycles (Table 2).^10,11^

### Experimental Protocol

Participants completed two sessions, the first session included accelerometry and metabolic testing, whilst the second session was overground biomechanical running trials. To ensure accurate indirect calorimetry readings, participants refrained from eating for at least 2 h and from consuming caffeine for at least 12 h before the start of each metabolic testing session. These testing sessions began with a standardized warm-up protocol where participants ran on the treadmill in their own shoes for 4 minutes at a speed 2 km.hr^−1^ slower than the testing speed followed by 6 minutes at the prescribed testing speeds: 14 km.hr^−1^ and 16 km.hr^−1^ for females and males, respectively. Footwear conditions for the subsequent testing trials were randomised across the three test shoes to reduce any potential order effects.

#### Session 1

Post warm-up, the runners completed two sets of 6-minutes of treadmill running in each footwear condition at 16 km.hr^−1^ (for males) or 14 km.hr^−1^ (for females). Specifically, each condition was repeated twice in a randomized, counterbalanced mirrored design, resulting in a total of six trials per participant. During the test, metabolic data were collected using indirect calorimetry (Quark CPET, COSMED, Rome, Italy), which was calibrated before each session according to manufacturer instructions. To ensure oxygen-consumption (VO₂) steady-state measurements, the speed selected was slower than the individual lactate threshold of each participant (further confirmed during the test by respiratory exchange ratios being below 1.0 during the whole running bout for all participants at each speed). Heart rate data were collected using the COROS heart rate strap positioned on participants’ upper arm in the space between the outer bicep and deltoid. Biomechanical data were collected during these trials with a commercial accelerometer (Runeasi, NV, Leuven, Belgium) placed on the lower back (between S2 and L5) of the participant.^12^ Rating of perceived exertion (RPE) scores were recorded within the last 30 seconds of each 6-minute trial, using the modified Borg scale from 0-10.

#### Session 2

Subsequent overground running trials for the determination of biomechanical factors were conducted on a 60-m indoor synthetic indoor sports floor. Participants completed 6 clean trials in the different footwear conditions at 4.44 m·s^−1^ (16 km.h^−1^, males) and 3.8 m·s^−1^ (14 km.h^−1^, females), in a randomised order. The running speed was measured using single-beam timing gates (Witty Microgate, Bolzano, Italy) placed at a 1-m height using a tripod and 4 m apart. Three-dimensional marker trajectories were captured using an 8-camera OptiTrack motion analysis system (NaturalPoint Inc., Corvallis, OR, USA), sampling at 240 Hz using a custom marker set. GRF data were collected using two 900 × 600mm force platforms (AMTI, Watertown, MA, USA), sampling at 1200Hz, synchronized with the motion capture system.

### Data Processing

#### Running Economy

VO₂ (mlO_2_·kg^−1^·min^−1^) values collected from the 2:30 to 5:30 mark of each running bout were averaged and designated as steady-state oxygen cost of transport for each trial. These were converted into Running Economy (mlO2·kg^−1^·km^−1^) values by accounting for the speeds at which the trials were performed. Running economy was further expressed in terms of the Energy Cost of running by accounting for the energy equivalent (in Joules) per unit of oxygen consumed. The respiratory exchange ratio was continuously monitored to ensure it remained below 1.00. Participant RPE data (Table 3) further indicated that they perceived the trials as “Easy” to “Moderate” at most.

**Table 3:**
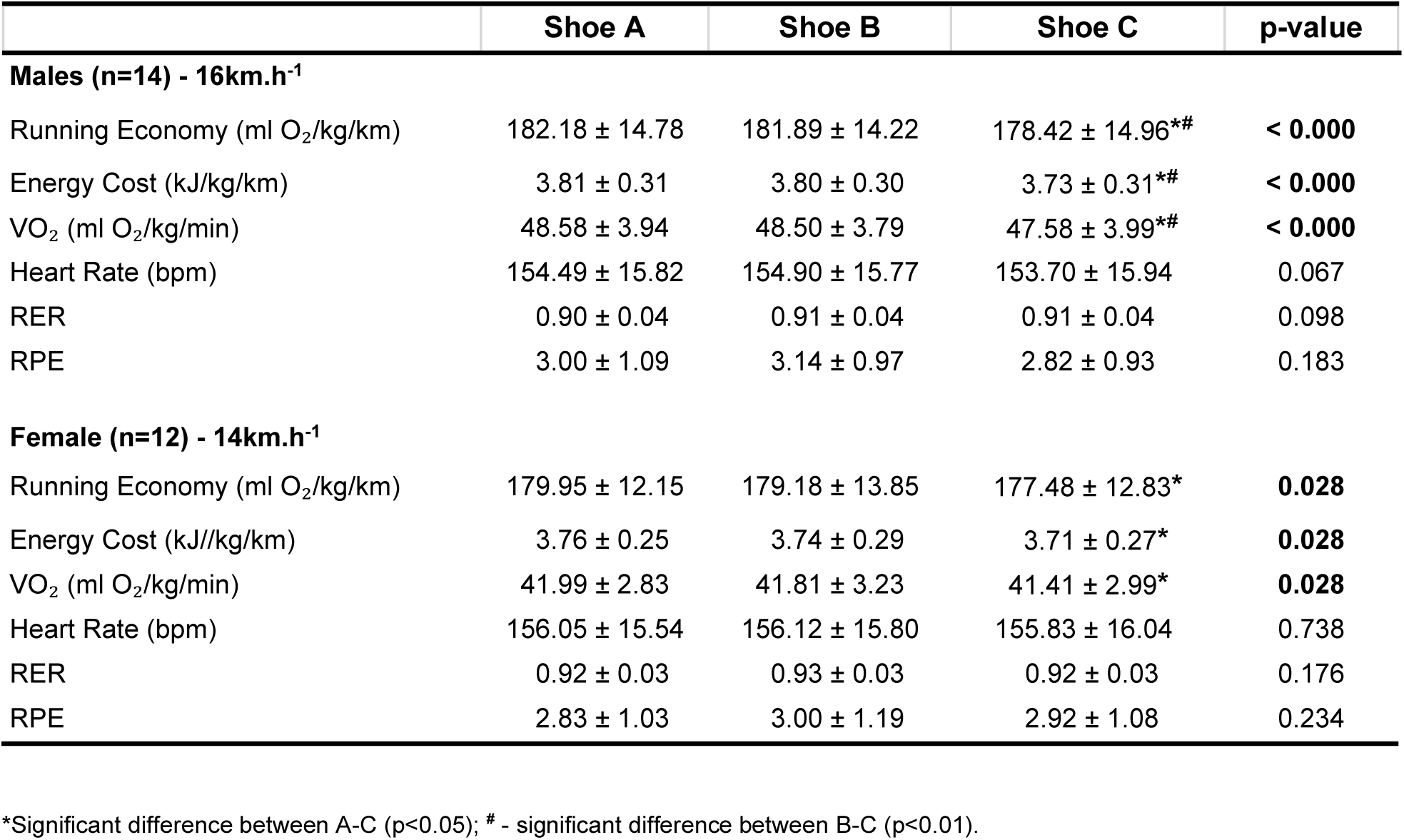
Physiological parameters of the male and female runners during the running economy test.

#### Biomechanical Variables

Lower-body biomechanics were modelled using the standard Plug-in Gait model, with kinematic and kinetic outputs calculated with all motion capture files saved in C3D format. All biomechanical data were imported into MATLAB (R2024a, The MathWorks Inc., Natick, MA, USA) for processing and analysis. Processed .C3D files were read into MATLAB using the open-source Biomechanical Toolkit.^13^ Each trial’s marker trajectories, joint kinematics, kinetics analog data and events were organized into a hierarchical structure indexed by participant, shoe condition and trial numbers. All subsequent processing, including event validation, time normalisation and computation of derived outcomes was performed in this MATLAB environment.

Marker trajectory and analog data were filtered using a low-pass fourth-order Butterworth filter with a cutoff frequency of 6 and 50 Hz, respectively. For each trial, 1 complete gait cycle was analyzed. The lower body Plug-In Gait model calculated 3-dimensional lower-extremity joint angles and net resultant joint moments using a Newton–Euler inverse dynamics approach using this data. Joint angles were described using the joint coordinate system. Three-dimensional joint moments were expressed as external moments normalized to body mass (N·m·kg^−1^). Hip, knee and ankle joint angles, joint velocity, moments and powers were extracted for each participant’s left limb and were averaged over the six trials.

Initial contact and toe-off events were auto-detected using a 5 N vertical ground reaction force threshold, while initial contact was designed as the instant the vertical force first exceeded 5 N, and toe-off as the instant the force subsequently dropped below this threshold. Stance phase was defined as the period between these two events. Trials were time-normalised to 101 data points (0-100% of the gait cycle) using cubic spline interpolation, with 0% defined as the initial contact of the left limb and 100% as the second contact of the left limb. For metrics that required stance phase specific time-normalisation (e.g. vertical ground reaction forces), the cubic spline method was applied to the period between the left initial contact and the subsequent left toe-off events, with the resulting waveforms re-normalised to 101 data points (0-100% of stance). Time-domain outcomes (e.g. time to pearl vertical ground reaction force) were computed directly from the raw non-normalised data using each trial’s native sampling rate, to preserve absolute temporal information.

Leg stiffness was calculated as described by Liew et al (2017).^15^ The leg vector definition was adjusted from hip-to-center-of-pressure to the hip-to-lateral-ankle, to isolate biological lower-limb compression from shoe-midsole compression. Leg compression was defined as the shortening of the leg vector from initial contact to midstance, and leg extension as the lengthening of the leg vector from midstance to toe-off.^16^ Joint quasi-stiffness was computed for the knee and ankle joints following the methods of Hamill et al. (2014).^17^ For each joint, the energy-absorption phase was defined as the period from initial contact to the first occurrence of peak knee flexion or peak ankle dorsiflexion within the stance phase for the knee and ankle, respectively. Quasi-stiffness was taken as the slope of a linear least-squares regression of sagittal-plane joint moment on joint angle within this window, multiplied by body mass to express it in absolute units (Nm·deg^−1^).

### Statistical Analysis

All statistical analyses were performed using Prism 11 (Graphpad Software, MA, USA). Data were screened for normality using the Shapiro-Wilk test prior to analysis. Male and female runners were examined as independent groups at running speeds matched to their respective competitive training intensities. No between sex statistical comparisons were conducted.

Differences in physiological outcomes across shoe conditions were examined using a one-way repeated measures ANOVA, appropriate for the within-subject crossover design in which each participant completed all shoe conditions. Prior to interpreting ANOVA results, the sphericity assumption was assessed using Mauchly’s Test. Where sphericity was violated (p<0.05), the Greenhouse-Geisser correction was applied to reduce the risk of Type I error. Where a significant main effect was found, Bonferroni-corrected pairwise post-hoc comparisons were conducted to identify specific differences between shoe conditions. Practical significance was assessed using Cohen’s d effect sizes, interpreted as small (0.2), medium (0.5), and large (≥ 0.8) (Cohen, 1992).^18^ Mixed-effects ANOVAs with Greenhouse-Geisser correction were used for the biomechanics and spatio-temporal data. A significance level of p<0.05 was applied to all analyses.

## Results

For the female participants, significant differences were found in RE, energy cost, and VO₂ between shoes (p=0.028), where Shoe C was 2.5 ml.kg^−1^.km^−1^ (∼1.4%) lower than Shoe A (p=0.004). Similar findings were noted in the male participants, specifically between Shoe A-C and Shoe B-C (p<0.000) (Table 3). Shoe C was approximately 3.8 ml.kg^−1^.km^−1^ lower than Shoe A (∼2.1%) (p=0.004) and 3.5 ml O₂/kg/km lower than Shoe B (∼1.9%) (p=0.002). Heart rate and RER were similar between all footwear conditions for both sexes (Figure 1,Table 3).

When observing the individual running economy responses across the different footwear, 5 (3 male, 2 female) of the 26 runners were less economical in Shoe C than Shoe A, 3 (1 male, 2 female) of the 26 runners were less economical in Shoe C than Shoe B, 15 (8 male, 7 female) of the 26 runners were less economical in Shoe B than Shoe A (Figure 2, Table 3).

**Figure 2:**
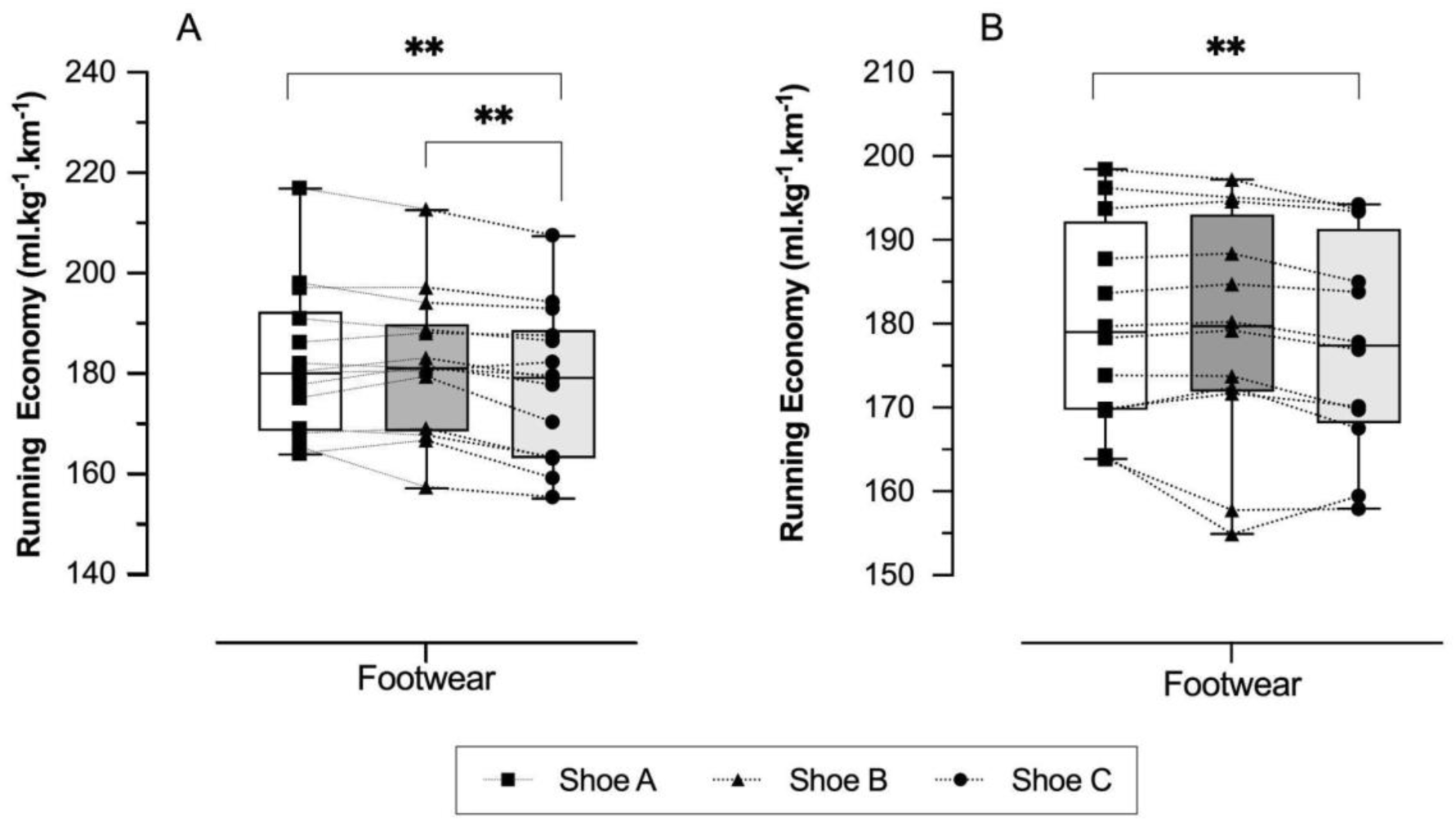
Individual running economy plots for both male (A) and female (B) runners across the three footwear conditions. ** - significant difference (p < 0.01)

**Figure 3:**
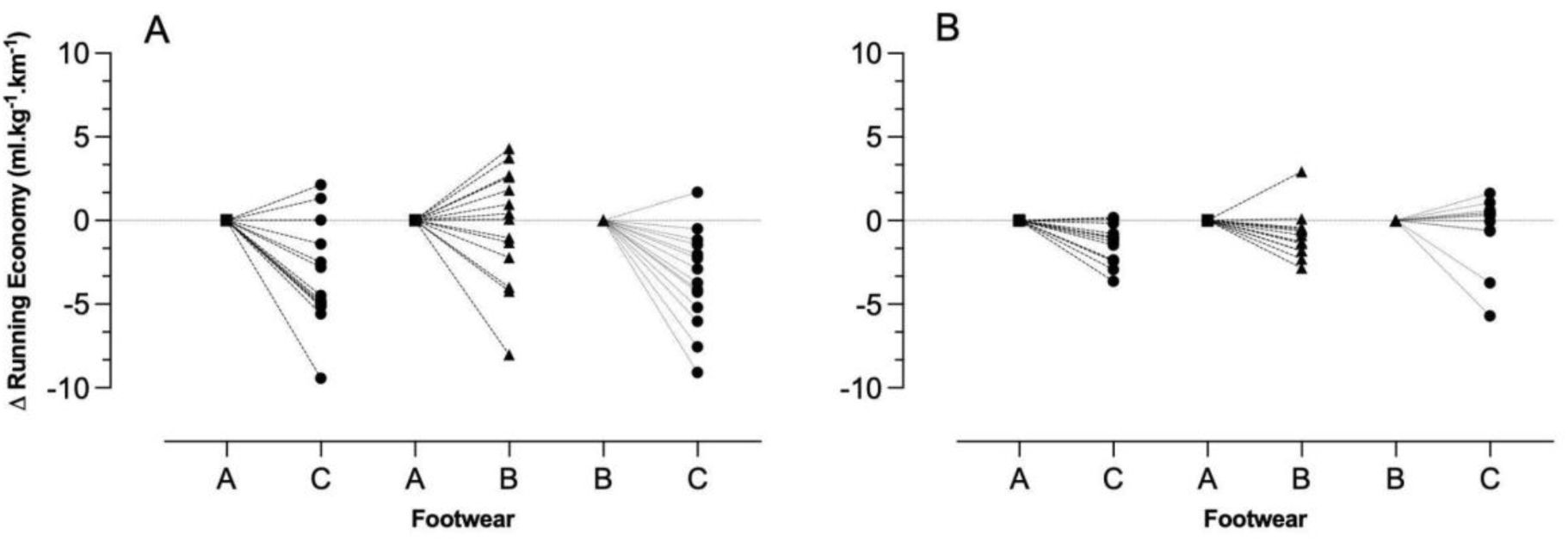
Individual changes in running economy for both male (A) and female (B) runners across the three footwear conditions.

Impact magnitude differed between footwear conditions in both females (p=0.003) and males (p=0.001), with Shoe A exhibiting lower impact magnitudes than Shoes B (p=0.014) and C (p=0.013) in both sexes. No differences were noted between Shoe B and C.

Further footwear effects on spatiotemporal variables were sex specific; males demonstrated longer impact duration in Shoe A than Shoes B and C (p=0.039), and higher cadence in Shoe A than Shoe C (p=0.031). Whereas females exhibited shorter ground contact times than both Shoe B (p=0.003) and Shoe C (p=0.001), with a higher cadence in Shoe A than Shoes B (p=0.034) and C (p=0.007; overall p=0.002). Furthermore, females showed a greater flight ratio between Shoe A and Shoe B (p=0.009) and Shoe C (p=0.007).

With regards to discrete kinematic and kinetic features, Ankle quasi-stiffness was found to be higher in Shoe A compared to Shoe C in male (p=0.043) and female runners (p=0.010). In the female runners, ankle quasi-stiffness in Shoe A was also higher than Shoe B (p=0.010).

Vertical ground reaction forces in female runners differed between Shoe A and B over 91–97% of stance (*F*\*=7.492, p<0.01) (Table 7, Figure 4). Shoe B exhibited less ankle dorsiflexion than Shoe A and C between 5-70% of stance (*F=6.377, p<0.0001) and Shoe A exhibited lower ankle angular velocity when compared to Shoe C over 25 - 38% of stance (*F=6.855, p<0.01) (Table 7, Figure 5). In the knee, greater angular velocity between 72-76% of stance was found in Shoe C when compared to Shoe B (*F=6.559, p<0.01), whereas between 81-90% of stance Shoe C was higher than Shoe A (*F=6.599, p<0.01). Knee moments were greater in Shoe A when compared to Shoe B during 78 - 96% of stance (*F=6.739, p<0.0001). In the hip, angular velocity was lower in Shoe A compared to Shoe B over 84-100% of stance (*F=6.543, p<0.0001). Whereas Shoe A presented greater moments 10% of stance when compared to Shoe B (*F=6.673, p<0.05).

**Figure 4:**
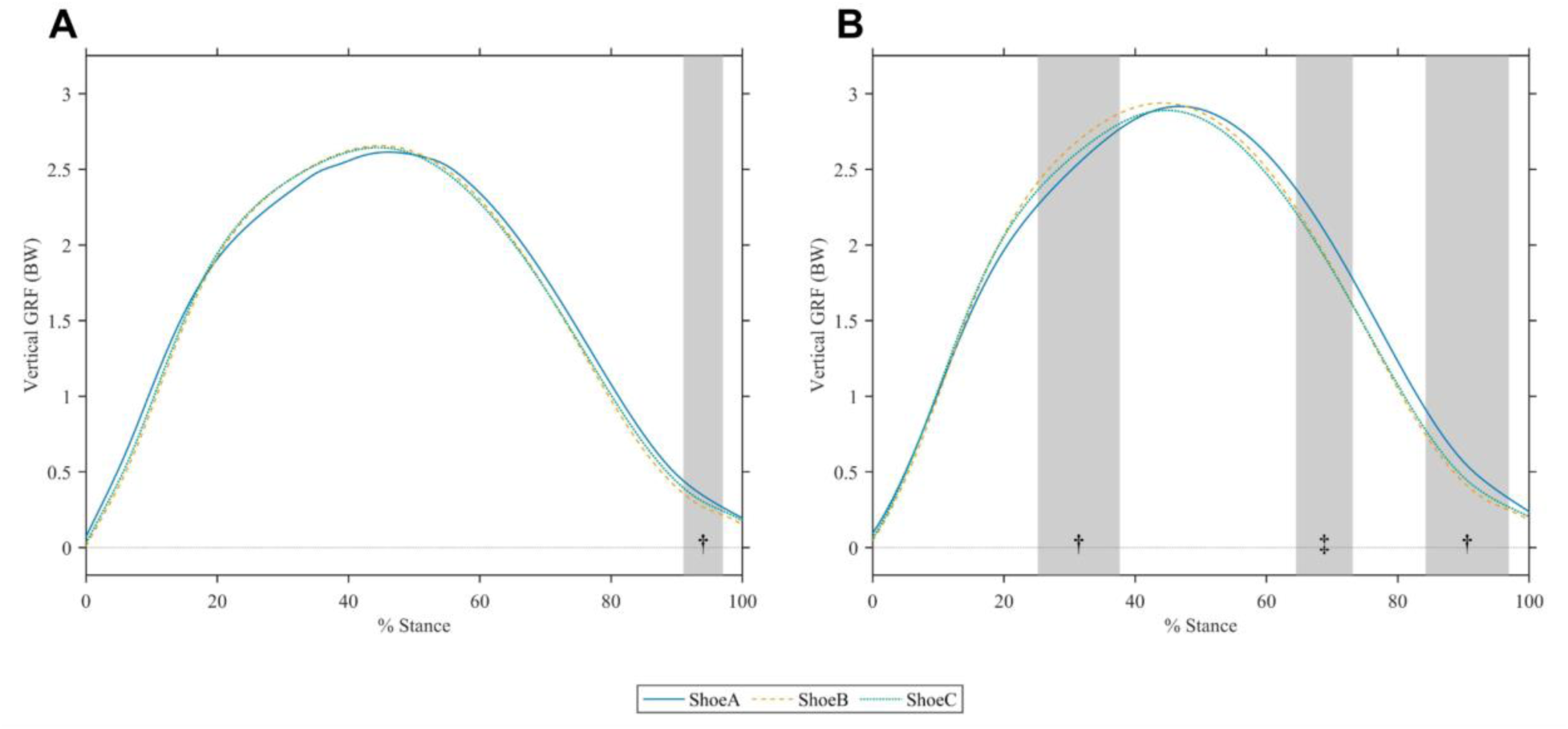
Mean vertical ground reaction force over 100% of stance in (A) female and (B) male runners in the different footwear conditions. Significant differences between conditions (grey shaded bands, ‡ - between Shoe A and B, ✝ - between Shoe A and C)

**Figure 5:**
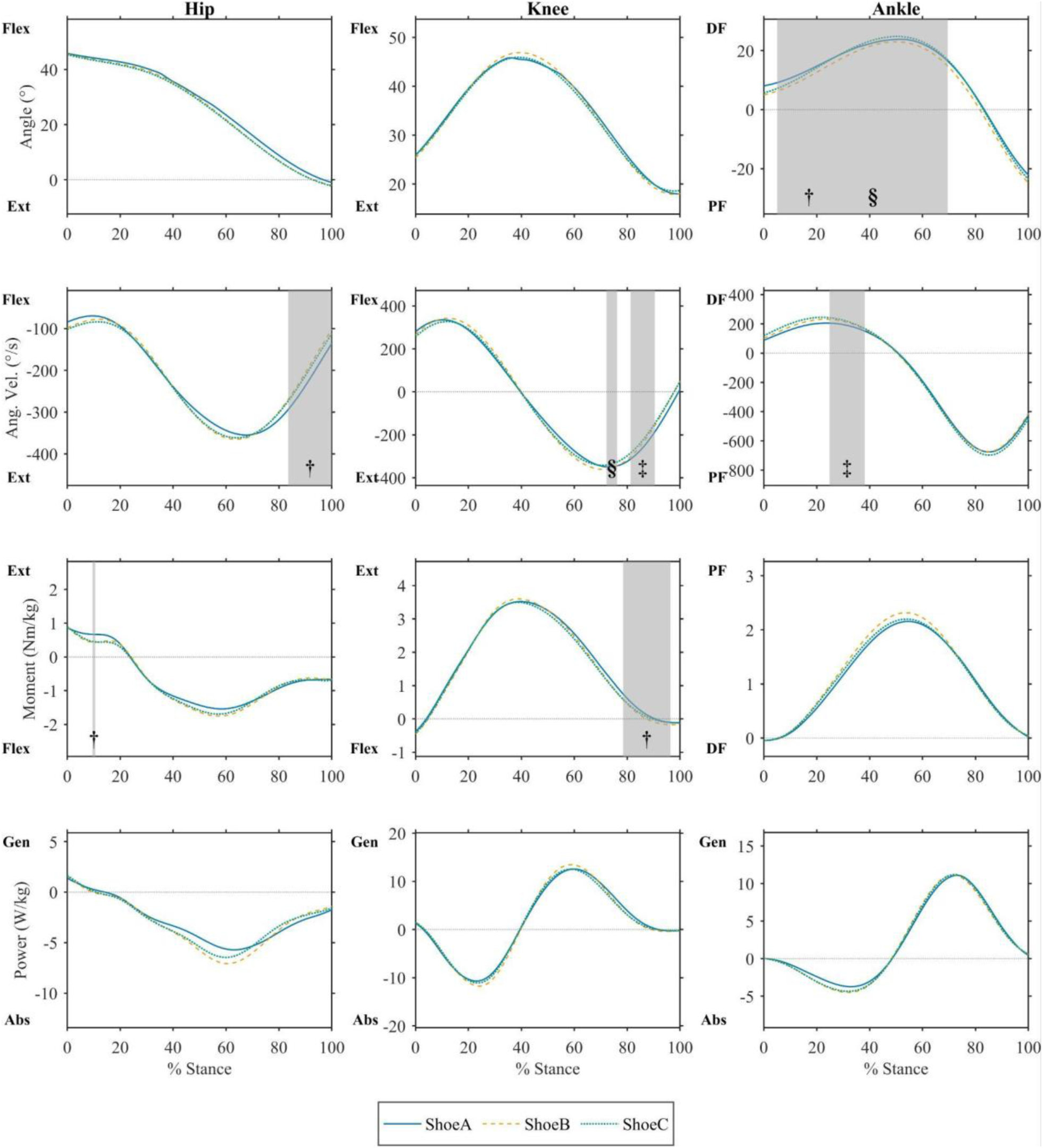
Kinematic and kinetics data over 100% of stance in the female runners (n=12). Significant differences between conditions (grey shaded band, ‡ - between Shoe A and B, ✝ - between Shoe A and C,§ - between Shoe B and C). Flex-Flexion; Ext- Extension; DF- Dorsiflexion; PF- Plantar Flexion; Gen- Generation; Abs-Absorption

In males, the differences in the ground reaction forces were found between Shoe A and B during 25 - 38% and 85 - 99% of stance (*F=7.190, p<0.001) (Table 7, Figure 4). Ground reaction forces differences were also found between Shoe A and C at 64 - 73% of stance (*F=7.190, p<0.01). Shoe C exhibited greater ankle dorsiflexion than Shoe B between 40 - 66% of stance (*F=6.007, p=0.001) and higher ankle angular velocity was found in Shoe B and C when compared to Shoe A during 28 - 36% of stance (*F=6.853, p<0.01) (Table 7, Figure 6). In the knee, greater angular velocity between 83-95% of stance was found in Shoe C when compared to Shoe A (*F=6.823, p<0.01). Knee moments were greater in Shoe A when compared to Shoe B and Shoe C during 71-92% of stance (*F=6.740, p<0.001). Similarly, knee power was higher in Shoe A compared to Shoe C over 73-89% of stance (*F=7.499, p<0.001). In the hip, angular velocity was lower in Shoe A compared to Shoe C over 87-100% of stance (*F=6.489, p<0.01). Whereas Shoe A experienced greater moments over 6-11% of stance when compared to Shoe B (*F=6.820, p=0.012).

**Figure 6:**
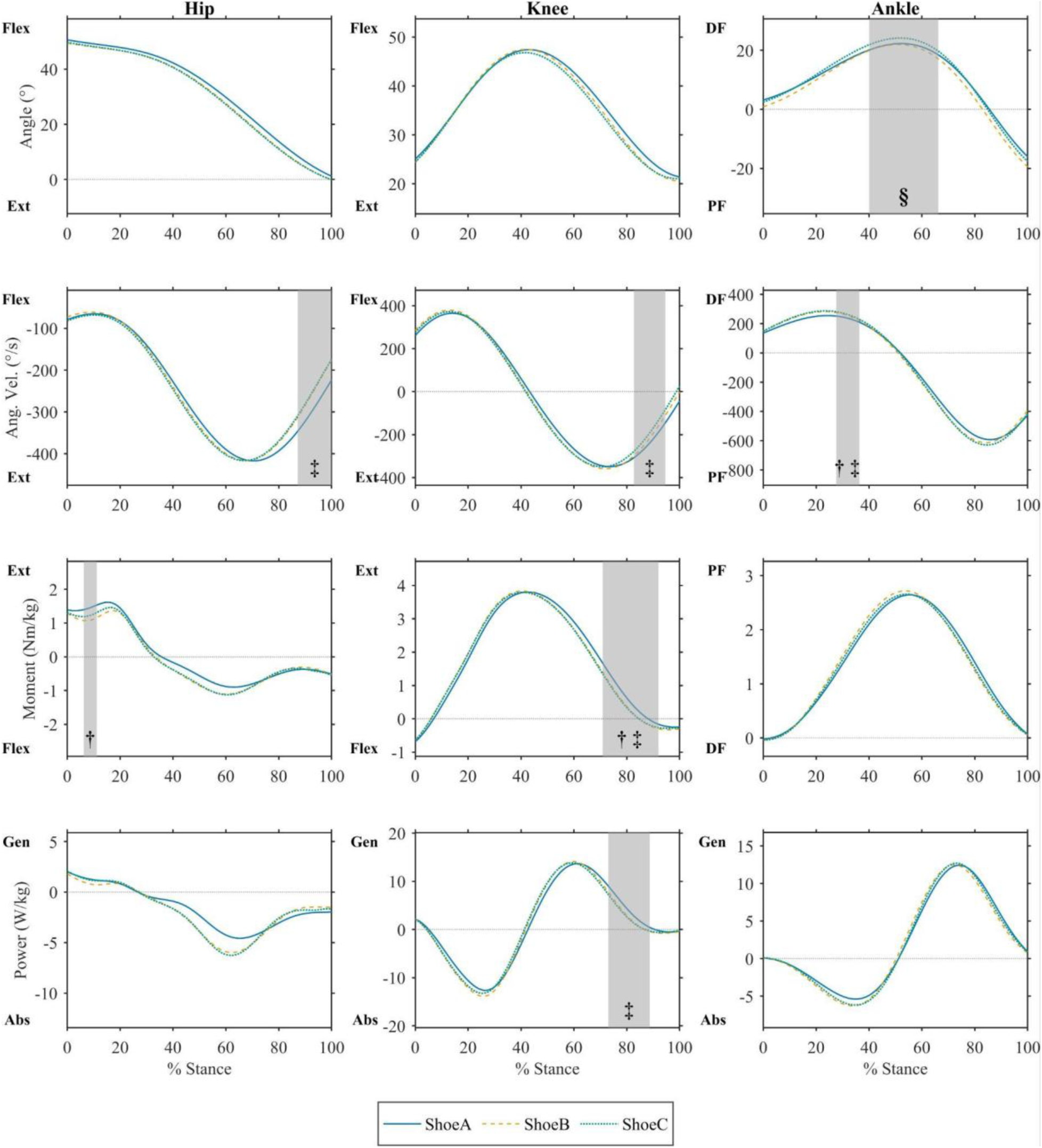
Kinematic and kinetics data over 100% of stance in the male runners (n=12). Significant differences between conditions (grey shaded band, ‡ - between Shoe A and B, ✝ - between Shoe A and C,§ - between Shoe B and C). Flex-Flexion; Ext- Extension; DF- Dorsiflexion; PF- Plantar Flexion; Gen- Generation; Abs-Absorption

## Discussion

The primary intention of this study was to improve our understanding of how well-trained runners respond to AFTs. We sought to understand whether males and females would elicit different physiological and biomechanical responses to AFTs when running at fast, metabolically stable sub-maximal running velocities. This study is noteworthy because of the calibre of runners recruited, who were capable of maintaining submaximal running speeds of 16-km.h^−1^ (3:75 min.km^−1^) for male runners and 14-km.h^−1^ (4:17 min.km^−1^) for female runners, at an average RPE of 3.00 ± 1.09 and 2.83 ± 1.06 respectively. This was important as it provided insight into the biomechanical effects of differences AFTs tested at velocities closest to racing pace than previously been investigated. Previous studies have compared AFT effects in cohorts running at the same relative intensities (e.g. 70% of VO₂_max_), and at slower running velocities. Therefore, it remains unclear whether any notable biomechanical changes are attributed to the workload (i.e. running velocity) or to the footwear intervention (i.e. shoes).^9^

The first important finding of this study was that despite different running velocity, male and female runners demonstrated similar physiological and biomechanical responses across the three different AFT conditions. Running economy was most efficient in Shoe C when compared to both Shoe A (1.4%, females and 2.1%, males) and B (0.9% females and 1.94%, males). Interestingly, improvements were consistent over the entire cohort in Shoe C when compared to Shoe A and B, and not as disparate as in previous studies,^19,20^ suggesting a general beneficial response by majority of the runners, however female runners showed smaller improvements in RE than the male runners in the different AFTs. Collectively, 73% (19/26) responded in Shoe C, with 78.6% males (11/14) and 66.7% females (8/12). The lower running velocity and body mass of the female runners may not have been sufficient to fully utilise the properties of the AFT and maximise its potential benefits.

Interestingly, the lightest AFT condition, Shoe A, demonstrated the lowest RE. This finding challenges the commonly cited assumption that a 100g reduction in mass is associated with a 1% improvement in running economy.^21^Franz and Kram (2012) achieved this relationship in a barefoot condition with added “weight” on a sock attached onto the ventral aspect of the foot.^21^ When assessing AFT midsole mass contribution, it is important to acknowledge not only the mass but also the contribution of this mass in providing an economical benefit. Specifically, mass with compliant properties added on the ventral aspect of the foot would be more useful than mass added on the dorsal aspect of the foot. Some studies have found that greater footwear midsole compliance controlling for other common metrics such as mass and bending stiffness resulted in improved running economy.^16^ It appears that the relative contribution of footwear midsole properties concomitantly with the associated mass require a fine balance to exert an improvement in running economy.

Mechanistically, previous research has also found that greater midsole compliance is associated with greater leg stiffness and peak vertical ground reaction forces.^16,22^ Differences in these variables were not observed between shoe conditions in the present study. Instead, we found greater impact magnitude in Shoe C when compared to Shoe A and B (Table 4). In addition, the higher forces at early stance between Shoe A-B and lower forces in late-terminal stance in Shoe A-B and A-C. Together these findings point to Shoe A exhibiting a more different loading pattern and impact manifestation to both Shoe B and C. This might indicate the performance benefits of high forces experienced during high running velocities are able to be utilised by the body through a more efficient coordinated spring-like behaviour.^14^ The impact magnitude reported here was derived from the sacral accelerometer, which serves as a proxy for the runner’s centre of mass that is indicative of the body’s response to the external workload (running speed).^12^ At lower running speeds, the legs are generally thought to actively dampen the impact forces through biomechanical mechanisms. However, at higher workloads, well-trained runners may be better able to effectively accept and utilise this impact energy with the midsole compliance acting to reduce the need for active biomechanical dampening of the leg.^23^

**Table 4:**
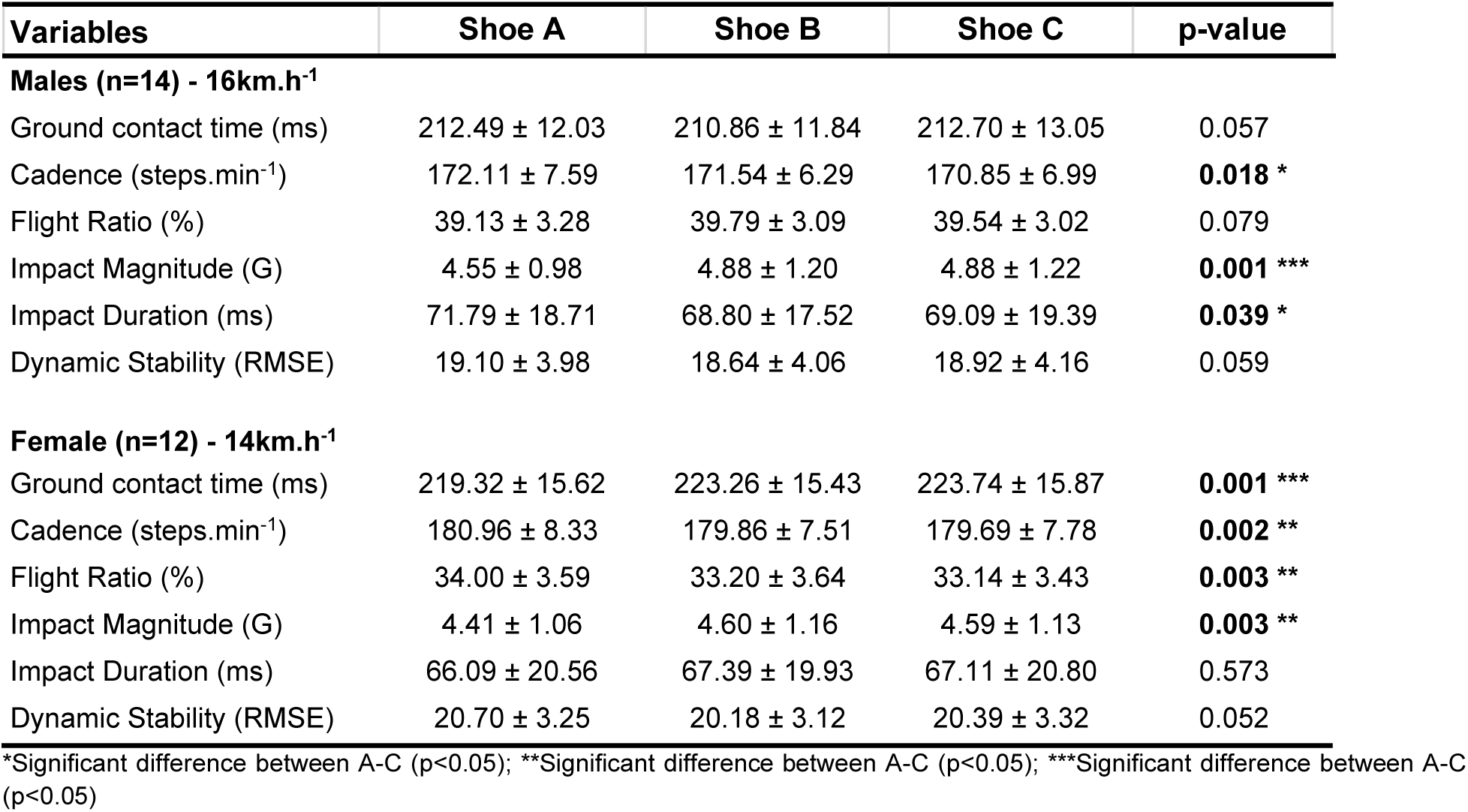
Spatiotemporal and biomechanical sensor metrics for male and female runners.

| Variables | Shoe A | Shoe B | Shoe C | p-value |
| --- | --- | --- | --- | --- |
| **Males (n=14) - 16km.h<sup>-1</sup>** |  |  |  |  |
| Ground contact time (ms) | 212.49 ± 12.03 | 210.86 ± 11.84 | 212.70 ± 13.05 | 0.057 |
| Cadence (steps.min<sup>-1</sup>) | 172.11 ± 7.59 | 171.54 ± 6.29 | 170.85 ± 6.99 | **0.018 \*** |
| Flight Ratio (%) | 39.13 ± 3.28 | 39.79 ± 3.09 | 39.54 ± 3.02 | 0.079 |
| Impact Magnitude (G) | 4.55 ± 0.98 | 4.88 ± 1.20 | 4.88 ± 1.22 | **0.001 \*\*\*** |
| Impact Duration (ms) | 71.79 ± 18.71 | 68.80 ± 17.52 | 69.09 ± 19.39 | **0.039 \*** |
| Dynamic Stability (RMSE) | 19.10 ± 3.98 | 18.64 ± 4.06 | 18.92 ± 4.16 | 0.059 |
| **Female (n=12) - 14km.h<sup>-1</sup>** |  |  |  |  |
| Ground contact time (ms) | 219.32 ± 15.62 | 223.26 ± 15.43 | 223.74 ± 15.87 | **0.001 \*\*\*** |
| Cadence (steps.min<sup>-1</sup>) | 180.96 ± 8.33 | 179.86 ± 7.51 | 179.69 ± 7.78 | **0.002 \*\*** |
| Flight Ratio (%) | 34.00 ± 3.59 | 33.20 ± 3.64 | 33.14 ± 3.43 | **0.003 \*\*** |
| Impact Magnitude (G) | 4.41 ± 1.06 | 4.60 ± 1.16 | 4.59 ± 1.13 | **0.003 \*\*** |
| Impact Duration (ms) | 66.09 ± 20.56 | 67.39 ± 19.93 | 67.11 ± 20.80 | 0.573 |
| Dynamic Stability (RMSE) | 20.70 ± 3.25 | 20.18 ± 3.12 | 20.39 ± 3.32 | 0.052 |

\*Significant difference between A-C ( $p<0.05$ ); \*\*Significant difference between A-C ( $p<0.05$ ); \*\*\*Significant difference between A-C ( $p<0.05$ )

**Table 5:** Discrete kinematic and kinetic variables for both male and female runners.

| Variable | Shoe A | Shoe B | Shoe C | p-value |
| --- | --- | --- | --- | --- |
| **Males (n=12)** |  |  |  |  |
| Leg Stiffness (N.mm<sup>-1</sup>) | 34.54 ± 16.43 | 32.42 ± 10.93 | 34.19 ± 14.56 | 0.369 |
| Leg compression (mm) | 61.02 ± 16.36 | 62.35 ± 14.97 | 59.85 ± 15.57 | 0.485 |
| Leg extension (mm) | 66.48 ± 12.94 | 68.63 ± 10.31 | 65.96 ± 10.98 | 0.143 |
| Knee Quasi-Stiffness (Nm.deg<sup>-1</sup>) | 13.71 ± 2.87 | 13.07 ± 2.43 | 13.25 ± 2.46 | 0.486 |
| Ankle Quasi-Stiffness (Nm.deg<sup>-1</sup>) | 9.63 ± 3.50 | 9.03 ± 2.73 | 8.67 ± 2.87 | **0.043\*** |
| **Females (n=12)** |  |  |  |  |
| Leg Stiffness (N.mm<sup>-1</sup>) | 29.52 ± 7.86 | 28.04 ± 8.58 | 30.79 ± 11.56 | 0.239 |
| Leg compression (mm) | 50.26 ± 14.67 | 54.05 ± 17.69 | 49.93 ± 16.39 | 0.065 |
| Leg extension (mm) | 60.70 ± 12.79 | 63.42 ± 11.22 | 61.47 ± 9.46 | 0.181 |
| Knee Quasi-Stiffness (Nm.deg<sup>-1</sup>) | 10.86 ± 2.31 | 10.661 ± 2.77 | 11.249 ± 3.52 | 0.551 |
| Ankle Quasi-Stiffness (Nm.deg<sup>-1</sup>) | 7.86 ± 2.49 | 7.39 ± 2.37 | 6.65 ± 2.07 | **0.010\$\*** |

\*- significant difference between A-C (p<0.05); # - significant difference between B-C (p<0.01); \$- significant difference between A-B (p<0.05).

**Table 7:** Summary of post-hoc Statistical Parametric Mapping results across biomechanical variables.

|  | Variable | Critical threshold exceeded (% of stance) | Critical threshold (*F) | Supra-threshold p-value | Footwear comparison |
| --- | --- | --- | --- | --- | --- |
| **Female** |  |  |  |  |  |
| *Hip* | Angular Velocities | 84-100 | 6.54 | $< 0.001$ | A-B |
| | Moments | 10-10 | 6.67 | $< 0.05$ | A-B |
| *Knee* | Angular Velocities | 72-76 | 6.60 | $< 0.01$ | B-C |
| | | 81-90 | | $< 0.01$ | A-C |
| | Moments | 78-96 | 6.74 | $< 0.001$ | A-B |
| *Ankle* | Angles | 5-70 | 6.38 | $< 0.001$ | A-B, B-C |
| | Angular Velocities | 25-38 | 6.86 | $< 0.01$ | A-C |
| *Ground reaction forces* | Vertical | 91-97 | 7.49 | $< 0.01$ | A-B |
| **Male** |  |  |  |  |  |
| *Hip* | Angular Velocities | 87-100 | 6.49 | $< 0.01$ | A-C |
| | Moments | 6-11 | 6.82 | $p = 0.012$ | A-B |
| *Knee* | Angular Velocities | 83-95 | 6.82 | $< 0.01$ | A-C |
| | Moments | 71-92 | 6.74 | $< 0.001$ | A-B, A-C |
| | Powers | 73-89 | 7.50 | $< 0.001$ | A-C |
| *Ankle* | Angles | 40-66 | 6.01 | $p = 0.001$ | B-C |
| | Angular Velocities | 28-36 | 6.85 | $< 0.01$ | A-B, A-C |
| *Ground Reaction forces* | Vertical | 25-38 | 7.19 | $< 0.001$ | A-B |
| | | 64-73 | | $< 0.01$ | A-C |
| | | 84-97 | | $< 0.001$ | A-B |

Additionally on a joint level, ankle quasi-stiffness was lower in Shoe C than Shoe A in both female and male runners, and lower in Shoe B than Shoe A in the female runners only. This is an interesting observation, as previous studies found better running economy was correlated with lower ankle stiffness in runners with varying performance levels not controlling for footwear.^23,24^ This emphasises the important contribution of the ankle joint to the metabolic cost of running. The ability of the ankle to rotate faster during the early stance (during loading) appears to be indicative of metabolic optimality in the musculotendinous shank complex to produce the necessary forces.^25^

This is further detailed where Shoe C was found to exhibit a great plantarflexion velocity leading up to midstance with a subsequent greater ankle dorsiflexion magnitude during midstance despite no observed significant changes in joint kinetics. Suggesting an efficient neuromuscular gearing during the loading phase of gait. Leading on from this, during terminal stance (post-peak vGRF) both hip and knee angular velocity were greater in Shoe C. This appeared to be possibly facilitated by lower knee extension moment and positive power at the same phase of gait.

The capacity of the body to be able to maximise the joint angular velocities from ankle through to the hip may be exemplary of the lower longitudinal bending stiffness of Shoe C compared to Shoe A and B. The exploration of footwear gearing has been a subject of great interest over the past decades.^26–28^ Not all of these differences were found between the male and female runners, however, the similar patterns in the gait waveform and gross kinetics were found between the two groups and different running velocities. Perhaps the slower velocity did not elicit sufficiently large shifts in mechanics to adapt to the required workload.

Future research should explore detailed neuromuscular insights that might assist us to better understand the origins of the metabolic cost of these biomechanical adaptations, as it appears that the muscular work demands of running in AFTs do differ according to some of our findings. Also, these findings could lead to better understanding of AFT use for both the etiology of running injury and performance optimisation, that warrant further investigation. Ideally, we would like to perform overground running economy tests that would better reflect the overground biomechanical demands of competitive running. However, maintaining a constant high running velocity, while ensuring a consistent running surface across trials, presents methodological challenges.

## Conclusion

Males and female runners exhibited similar physiological and biomechanical responses to differing AFTs. The most distinct difference observed was that the female runners presented smaller improvements in running economy between AFTs, which could be attributed to the protocol running speed, body mass differences interacting with the properties of the AFTs. Notably, the large individual responses to the different AFTs were not observed in Shoe C, suggesting that the AFT properties do influence runners’ metabolic responses differently. Lastly, we noted the complexity of the biomechanical changes that resulted in the differences between AFTs, where greater vertical loading rate and impact magnitude kinetic metrics were found in the most efficient AFT alongside greater angular velocities of the ankle in early to mid-stance and the knee and hip in late to terminal stance. Concomitantly these observations suggest that the most efficient AFT enable these well-trained runners to be more spring-like through tolerating higher forces and faster angular velocities without greater demand on metabolic cost.

## Acknowledgements

Trevino Larry, Jordan Leondiris, Zabanguni Phakathi, Gian-Andri Baumann for their support and contributions towards this study.

